# Adult Neurogenesis in Peripheral Nervous System

**DOI:** 10.1101/2020.11.06.371880

**Authors:** Yisheng Liu, Xiaosong Gu

## Abstract

Although postnatal neurogenesis has been discovered in some regions of the peripheral nervous system (PNS), only indirect evidences indicated that some progenitors in the adult sciatic nerve and dorsal root ganglion (DRG) serve as a source of newly born sensory neurons. Here, we report the discovery of neurons and neuronal stem cells in the adult rat sciatic nerve. Lineage tracing detected a population of sciatic nerve neurons as progeny of adult neuronal stem cells. With the further finding of labeled DRG neurons in adult transgenic rats with local sciatic nerve staining, we propose a model of adult neurogenesis in the sciatic nerve in which neuronal stem cells in sciatic nerve mature as sensory neurons in adults along the sciatic nerve to DRG. This hypothesis provides a new way to understand sensory formation in adults. Those neuronal stem cells in the sciatic nerve may help to therapy of nerve trauma and disease in the future.

## Introduction

Neurogenesis occur in specific regions of the mammalian brain throughout life with critical roles in brain plasticity such as learning, memory and mood regulation (Berg et al., 2019). Postnatal peripheral nervous system neurogenesis has been discovered in mammalian parasympathetic ganglia of the head(Dyachuk et al., 2014; Espinosa-Medina et al., 2014) and the gut(Uesaka et al., 2015). This finding suggests that neurogenesis might also occur in the adult PNS, such as in the sciatic nerve and dorsal root ganglion (DRG).

However, confirming the in vivo existence of neuronal stem cells in these regions is challenging. In contrast to the extensive research of adult neurogenesis in the mammalian brain, we know very little about the adult neuronal progenitors in the PNS. Recent studies have revealed stem-like populations in DRG that displayed sphere-forming potential and multipotency in vitro, yet the in vivo presence of neuronal stem cells in the DRG has not been documented in any ultrastructural studies in adult mammals(Li et al., 2007; Nagoshi et al., 2008; Vidal et al., 2015). The sensory DRG neurons are derived from the thoracolumbar region of the trunk neural crest. The cervical region of those neural crest differentiate into large diameter neurons at first(Lawson and Biscoe, 1979). Late emigrating trunk neural crest give rise to boundary cap neural crest stem cells, a source of multipotent sensory specified stem cells(Radomska and Topilko, 2017). As a transient population, the embryonic neural crest quickly transfer from multipotent to restricted progenitors with limited capacity to self-renew before birth (Bronner and Simoes-Costa, 2016). In mammalian fetal and adult peripheral nerves and skin, neural crest derivatives give rise to multiple derivatives in vitro(Gresset et al., 2015; Morrison et al., 1999; Wong et al., 2006). This finding suggests that a subset of the neural crest population in the sciatic nerve and skin maintain multipotency after embryonic development. But the identity of precursors to adult sciatic nerve and DRG neurons and how they maintain their multipotency during development from embryonic to adult mammals are unknown.

One major obstacle to studying the adult neurogenesis in PNS is a lack of methods to the identification of neuronal stem cells in vivo. Many studies focus on cell isolation or in vitro culture of adult sciatic nerve and DRG because of the lack of a more specific in vivo tool(Baggiolini et al., 2015; Morrison et al., 1999). We established a sciatic nerve crush model in adult rats. By whole-mount staining and optical imaging of the crushed sciatic nerve tissue for stathmin 2 (Stmn2, or Scg10) (Shin et al., 2014), we observed neurons and neuronal stem cells in adult rat sciatic nerve. As an intermediate filament protein in neuroepithelial precursor cells, Nestin is considered a hallmark of neural stem/progenitor cells(Dubois et al., 2006; Hockfield and McKay, 1985; Lendahl et al., 1990; Wiese et al., 2004). We characterized neuronal stem cells labeled by the Nestin-CreER^T2^ rat line and Nestin-Cre rat line in the adult sciatic nerve but not in DRG with clonal lineage-tracing. In adult rats stained for the neuron marker Stmn2 and Peripherin(Escurat et al., 1990), the lineage-tracing neuronal stem cell and its progeny were temporality and spatiality distributed along the sciatic nerve from the dermal nerve ending to the DRG, suggesting that adult neurogenesis in the DRG does not occur in situ but, rather, new cells migrate along the sciatic nerve. During embryonic development, the neural crest migrate from the neural tube to the DRG as sensory neurons and to the sciatic nerve and dermal nerve ending as multipotent cells(Baggiolini et al., 2015; Gresset et al., 2015; Morrison et al., 1999). Our study provides a new perspective that those multipotent cells will mature in the sciatic nerve and migrate from sciatic nerve to DRG as sensory neurons in adult.

## Results

### The Adult Sciatic Nerve Contains Neuronal Cell Bodies

To enable us to image deep within PNS structures, we used a clearing reagent called ScaleS that renders the rat DRG and sciatic nerve transparent, but completely preserves fluorescent signals from labeled cells(Hama et al., 2015). Optical clearing of tissue allowed us to identify neuronal cell bodies in the sciatic nerve. Three days after creating a 1mm sciatic nerve lesion in twenty adult rats for 30s crush by forceps, we used an Stmn2 antibody to identify regeneration projections of the damaged DRG neurons. Surprisingly, we observed neuron-like cells at the distal end of six rats sciatic nerve, though 14 other rats did not contain these cells (Fig. 1a). As mature Schwann cells generate Schwann spheres and pigment cells in crushed sciatic nerve in vitro and in vivo, the existence of the neuron-like cells may have been induced by injury(Takagi et al., 2011). To assess whether the neuron-like cells exist in undamaged nerves, we used the same optical clearing method on intact sciatic nerves of adult rats. We found that the neuron-like cells were also present in intact sciatic nerves in five of 25 control rats(Fig.1b).

**Fig. 1.**
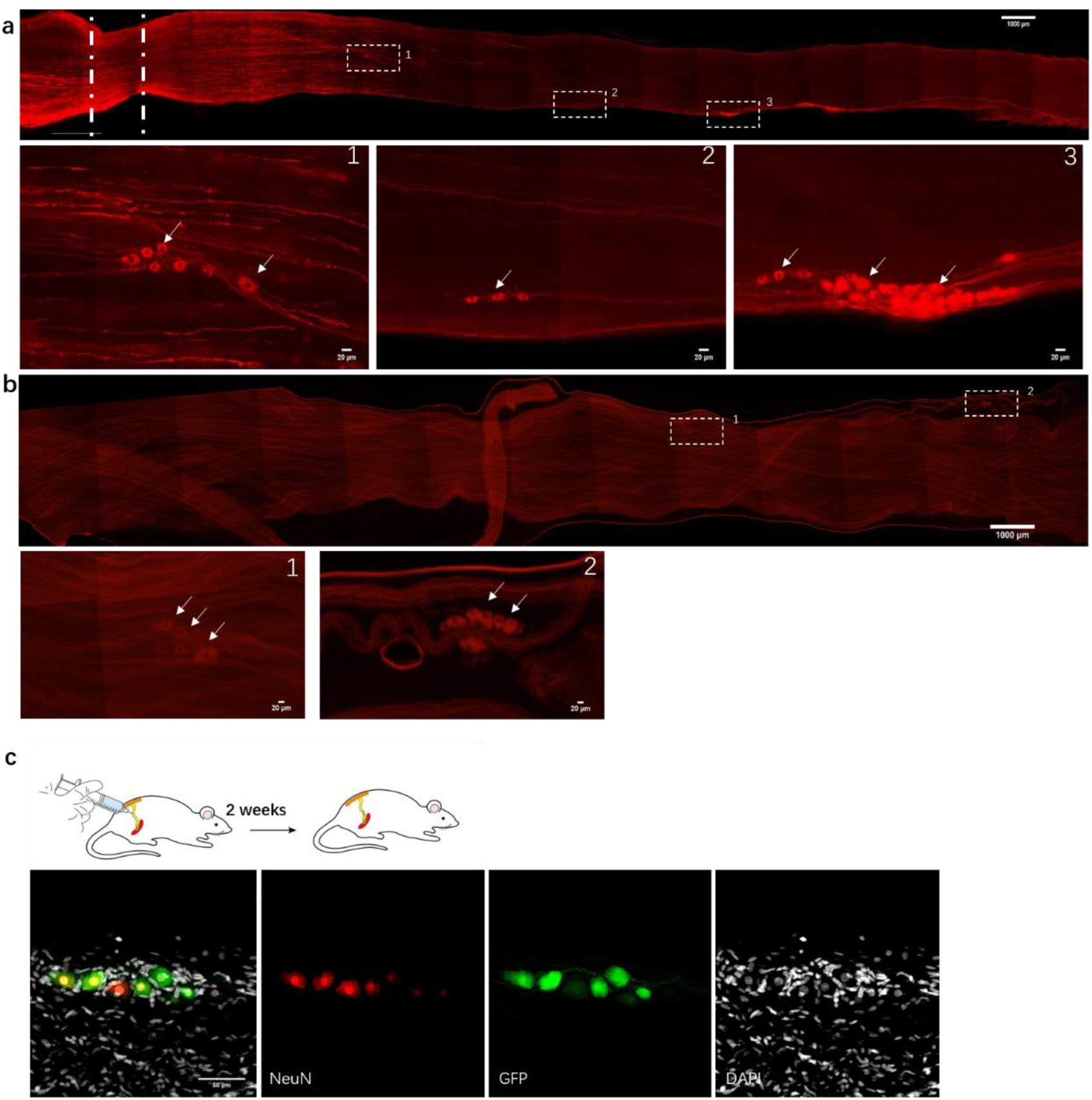
Different methods used to identify neurons in vivo in the adult rat sciatic nerve. **a**. Neuron-like cells in the crushed sciatic nerve of an adult rat. Lower panels show high-magnification images of the boxed regions in the upper panel. The sciatic nerve was hyalinized using ScaleS and stained for Stmn2 (red). The image shows neuron-like cells in the sciatic nerve. The white line indicates the crush site. (bar: upper =1000μm, lower=20μm) **b**. Neuron-like cells in the intact sciatic nerve of an adult rat. The nerve was hyalinized using ScaleS and stained for Stmn2 (red). The image shows neuron-like cells in the sciatic nerve. (bar: upper =1000μm, lower=20μm) **c**. Neuron-like cells labeled by AAV2/9 virus in vivo. NeuN and DAPI staining of virus-marked cells in vivo. Immunofluorescence staining for NeuN (red) and DAPI (gray) in GFP (green)-labeled cells in adult rat sciatic nerves 2 weeks after injection of AAV2/9 virus. (bar=50μm)

The unpredictable existence and location of those neuron-like cells in the sciatic nerve make those cells difficult to trace and identify. Delivery of adeno-associated virus (AAV) provides a noninvasive method for broad gene delivery to the nervous system(Foust et al., 2009). To label the neuron-like cells in vivo, we infected the sciatic nerves with engineered AAV2/9 and the hSYN and hEF1a promoter, to elicit stable expression of green fluorescent protein (GFP) in the cells. We detected the neuron projections in these infected neuron-like cells in vivo (Fig. 1c). Forty-six of 140 rats had these neuron-like cells in their sciatic nerves. Although only some of the neuron-like cells were stained, the expression of the neuron-specific marker NeuN(Mullen et al., 1992) indicated that part of those cells were indeed neurons (Fig. 1c). Together, this evidence indicates that there are neurons in adult rat sciatic nerve.

### Neuronal Stem Cells in Adult Sciatic Nerve

Because of the infrequent existence of the neurons in the sciatic nerves, we used an in vitro method to culture the sciatic nerve in a defined serum-free medium. We infected the adult rat sciatic nerve with AAV2/9 containing the hSYN and hEF1a promoter in vitro so that those cells stably expressed GFP, and observed labeled cells in sciatic nerves four to seven days later(Fig. 2a). For two weeks culture, sciatic nerves of 40 rats contained labeled cells while sciatic nerves of 170 rats were without. We then identified those cells by staining for peripherin, a marker of peripheral neurons (Fig. 2b).

**Fig. 2.**
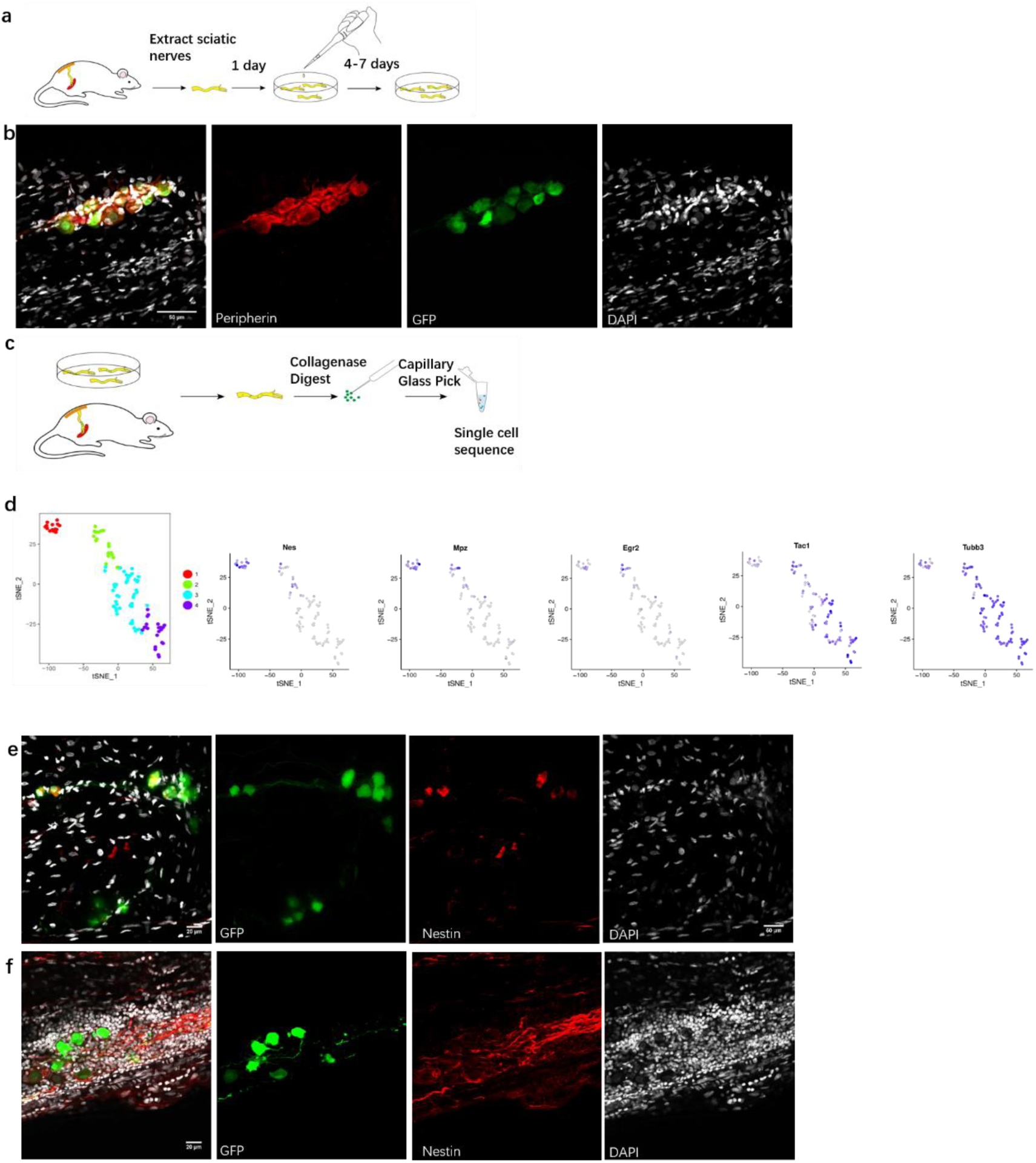
Neuronal stem cells in sciatic nerves of adult rats. **a**. Sciatic nerve culture model. The sciatic nerves of adult rats were cultured in a defined serum-free medium. **b**. Peripherin staining of virus-labeled cells in vitro. Immunofluorescence staining for peripherin (red) and DAPI (gray) in GFP (green)-labeled cells was performed in cultured adult rat sciatic nerves 2 weeks after injection of the AAV2/9 virus. (bar=50μm) **c**. Model of single-cell sequencing of virus-labeled neuron-like cells. **d**. t-SNE map representing the subcluster analysis of the 100 GFP+ cells that expressed high levels of Stmn2. The four colors represent four different clusters. Expression profile of neural crest differentiation related genes in the t-SNE map. Blue gradient represents the level of gene expression. **e**. Stem cell marker of neuron-like cells in vitro. Immunofluorescence staining for Nestin (red) and DAPI (gray) was performed in cultured adult rat sciatic nerves 2 weeks after injection of AAV2/9 virus in vitro. (bar=20μm) **f**. As for (e), but in vivo, using adult rat sciatic nerves 2 weeks after injection of AAV2/9 virus. (bar=20μm)

Next, to prospectively identify the labeled cells enables us to directly examine their properties at the molecular level. We conducted single-cell sequencing after 2 weeks of in vitro culturing(Fig.2c). 114 cells were dissected from GFP-positive sciatic nerves in vitro and in vivo (9 cells were from in vivo nerves). 10 DRG neurons were used as a positive control. Unsupervised clustering analysis assessed the separation of DRG neurons, Stmn2^+^ cells, and Stmn2^−^ cells(Blondel et al., 2008). Cells were separated into four clusters by further unsupervised clustering analysis. Cluster 1 and 2 cells showed high expression of transcripts for Egr2 (Krox20), nestin, and Mpz (protein 0), which are markers of migrating neural crest cells during the early fetal period(Dupin and Sommer, 2012). Cluster 3 and 4 cells showed high expression of Tac1 and Tubb3, which encode markers of mature neurons(Hokfelt et al., 2001; Jiang and Oblinger, 1992) (Fig.2d). As many cell-cycle genes were commonly expressed to varying degrees, this finding suggest that the labeled cells include neurons (some cells in cluster 4) and quiescent neuronal stem cells(some cells in cluster 1) that expressed low levels of cell-cycle genes, active neuronal stem cells(some cells in cluster 2,3) that expressed high levels of cell-cycle genes in the adult sciatic nerve.

We used a set of immunobiological markers and morphological criteria to identify and quantify the different cell types labeled by AAV2/9 in the sciatic nerve in vivo and in vitro. In both contexts, some of the cells exhibited radial glia-like morphology and expressed Nestin (Fig. 2e-f).

Together, these findings suggest that neuronal stem cells which we called multipotential sciatic nerve neural crest stem cells(snNCSCs) exist in the adult sciatic nerve in vitro and in vivo.

### Sciatic Nerve Neural Crest Stem Cells Differentiate Gradually in Adult Sciatic Nerve

To examine the cell fate of Nestin-positive cells in the adult rat sciatic nerve, we constructed a Nestin-CreER^T2^ rat for sequential observation(Dubois et al., 2006). We analyzed recombination in situ using reporter rats carrying an Tdtomato transgene whose expression was dependent on Cre-mediated recombination.

At different time points after tamoxifen and EdU intraperitoneal injection, We used stmn2 to identify and quantify cells labeled with Tdtomato in the sciatic nerves (Fig3a). The first two or three weeks, Tdtomato^+^ cells were not co-labeled with stmn2 but in the segments consisted of the stmn2^+^ cells. Most of those cells are set aside in quiescence without co-labeled with EdU(Fig3b). At four or more weeks, part of Tdtomato^+^ cells were co-labeled with stmn2 and EdU. Some of the Tdtomato^+^ stmn2^-^ cells and Tdtomato^-^ stmn2^+^ cells both co-labeled with EdU indicated those cells not only able to be differentiation but also self-renew (Fig3c). It consist with our single cell sequencing result and indicated most of those cells are quiescent neuronal stem cells and progenitors. For a long term tamoxifen injection, all the stmn2^+^ cells we detected are co-labeled with Tdtomato in 8 monthes old rats. This indicated those stmn2^+^ cells are derived from the Tdtomato^+^ cells.

**Fig. 3.**
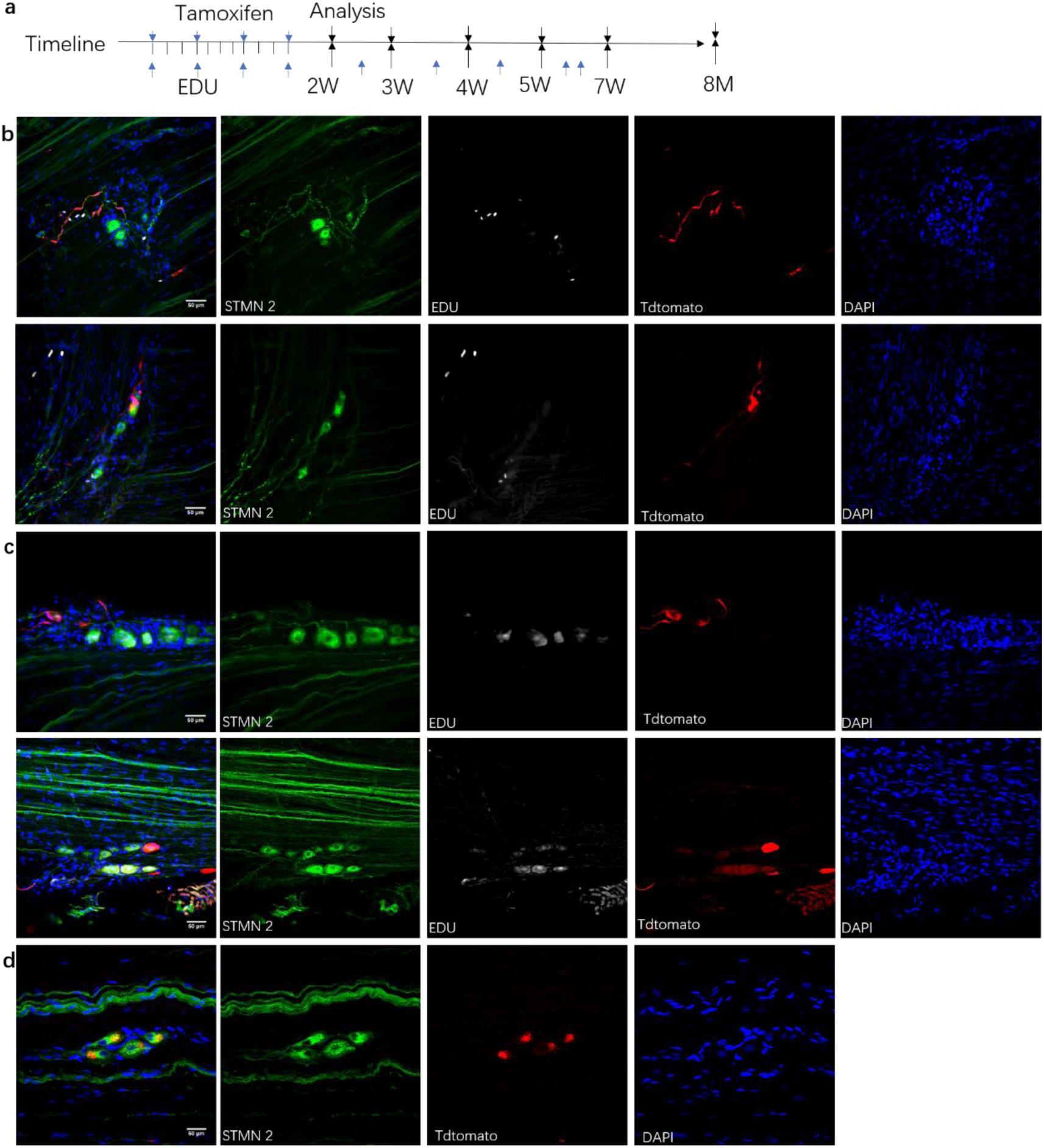
Neuronal stem cells gradually development in adult sciatic nerve. **a**. Adult Nestin-CreER^T2^::Tdtomato rats were given injections of tamoxifen and EdU at different time points for clonal lineage-tracing analysis. **b**. Confocal images of stmn2-labeled cells at 2 or 3 weeks after tamoxifen injection, but cells were not co-labeled with Tdtomato and EdU. (bar=50μm) **c**. Confocal images of stmn2-labeled cells at 4 or 5 weeks after tamoxifen injection co-labeled with Tdtomato and EdU. (bar=50μm) **d**. Confocal images of stmn2-labeled cells at 8 months after tamoxifen injection co-labeled with Tdtomato. (bar=50μm)

All those together, we concluded that the snNCSCs will set aside in quiescence and gradually differentiate into progenitors and neurons in the adult sciatic nerves in vivo.

### DRG Contains Neurons from Adult Sciatic Nerve

To investigate the in vivo differentiation potential and the final location of Nestin^+^ cells and their progeny, we utilized a transgenic approach in rat using the Nestin-Cre driver for labeling(Dubois et al., 2006) and performed mCherry staining of whole peripheral nerves and DRGs three weeks after tail vein injection of mCherry DIO (FLEx switch under Cre induction) AAV-PHP.S with hEF1a promoter virus in 12 nestin-cre^+/+^ rats. We dissected the L4 or L5 DRG and sciatic nerve of those rats. Surprisingly, NeuN staining revealed mCherry^+^ neurons in DRG of 3 rats (Fig. 4a). Next, we sacrificed rats 7 weeks after tail vein injection of AAV-PHP.S virus. Nineteen of 20 successfully injected rats contained mCherry^+^ neurons in their DRG(Fig. 4b).

**Fig. 4.**
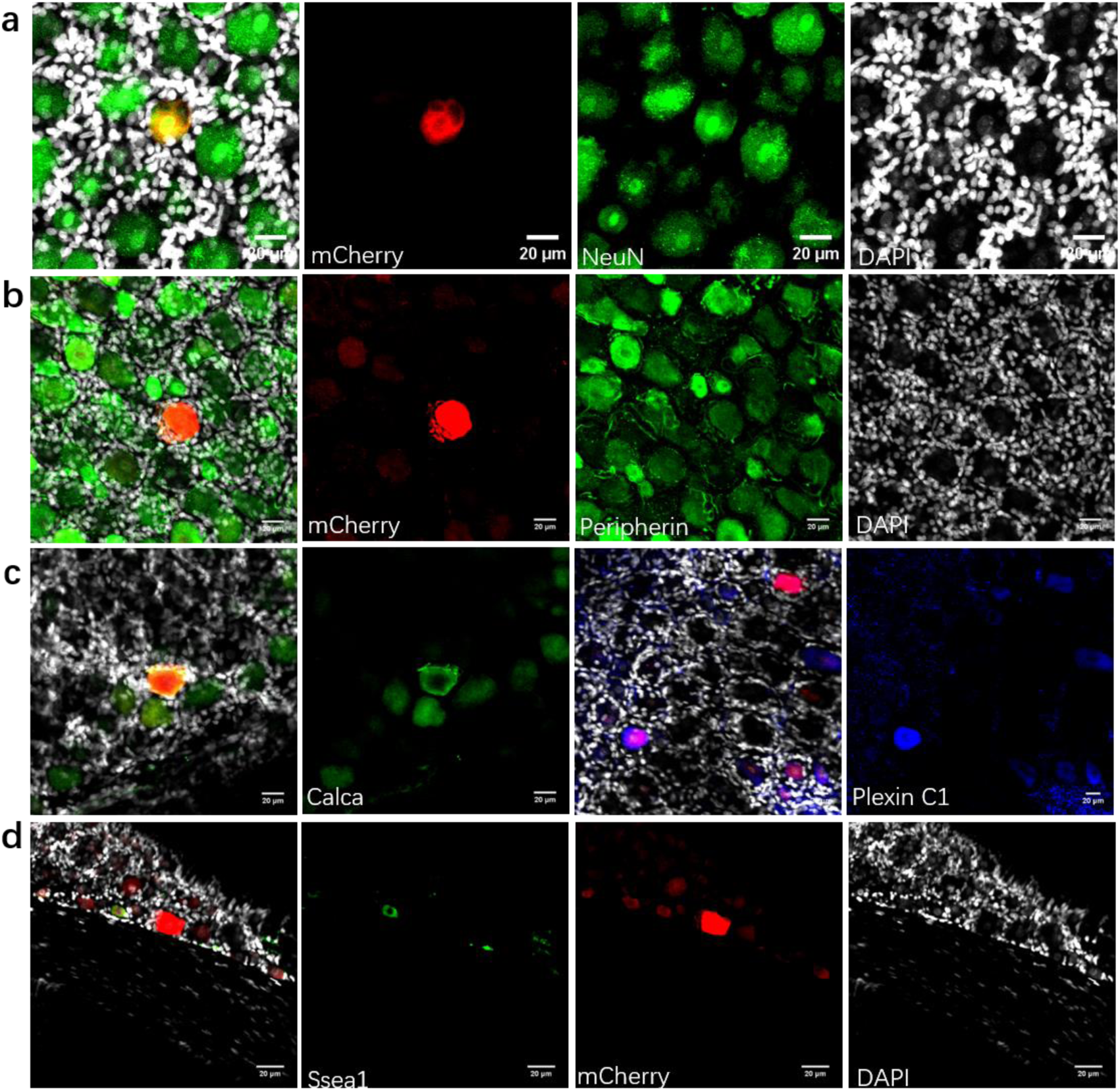
Distribution of newborn neurons from the sciatic nerve in the adult DRG. **a**. mCherry^+^ neurons in the DRG of Nestin-Cre^+/+^ rats subjected to tail vein viral injection. Immunofluorescence staining for NeuN (green) and DAPI (gray) was performed in mCherry-marked cells (red) of the adult Nestin-Cre^+/+^ rat DRG injected with AAV-PHP.S virus in vivo 3 weeks ago. (bar=20μm) **b**. As for (b), but for 7 weeks., using peripherin (green) and DAPI (gray) to stain DRG.(bar=20μm) **c**. mCherry-positive cells in the DRG. Immunofluorescence staining for DAPI (gray) and neuron-specific markers Calca (green) and Plexin C1 (blue) was performed in mCherry-labeled DRG cells (red) of the adult Nestin-Cre^+/+^ rat after AAV-PHP.S virus injection in vivo. (bar=20μm) **d**. Stem cell markers of viral-labeled cells in the DRG of Nestin-Cre^+/+^ rats. Immunofluorescence staining for Ssea1 (green) and DAPI (gray) was performed in mCherry-labeled DRG cells (red) of the adult Nestin-Cre^+/+^ rat after AAV-PHP.S virus injection. (bar=20μm)

To specifically assess the cell fate of the marked DRG cells, we injected the sciatic nerve of Nestin-Cre rats with hSYN-GFP DIO (FLEx switch under the induction of Cre) AAV-PHP.S virus, and sacrificed ten rats 3 weeks later. In 2 rats, GFP^+^ cells were observed in the DRG and sciatic nerve ipsilateral to injection with virus, whereas no GFP-positive cells were observed on the contralateral side. We further characterized traced cells with markers for different categories of mature DRG neurons, including peptidergic sensory neurons, such as Plexin C1 and Calca-positive neurons(Usoskin et al., 2015)(Fig. 4c). In addition, co-labeled with Ssea1(Sieber-Blum, 1989) but not Nestin or Egr2 indicated the limited differentiation potential of mCherry^+^ cells and newly born neurons in the DRG of mCherry DIO AAV-PHP.S-injected Nestin-Cre rats (Fig. 4d). Together, these findings indicate the possibility that neurons from the sciatic nerve migrate into the DRG as sensory neurons. The spatial distribution of the labeled neurons indicated a wave of sensory neuron migration from the sciatic nerve to the DRG in adult rats.

## Discussion

Although some subpopulations of NCSCs with high plasticity and sphere-forming capacity persist in the sciatic nerve and DRG during the late fetal stage to adulthood in vitro, it is technically challenging to demonstrate that post-migratory neural crest cells are maintain their multipotent properties and developmental plasticity in vivo(Bronner and Simoes-Costa, 2016; Parfejevs et al., 2018). Our discovery of neurons and neuronal stem cells in the adult mammalian sciatic nerve indicates that the nerve contains multipotent stem cells that differentiate into neurons. Our findings moreover indicate that sciatic nerve neurons will migrate into the DRG in adult.

As the existence and location of those cells are unpredictable for now in the adult sciatic nerve, we hypothesize that neural crest cells at the neural tube migrate to the sciatic nerve and dermis nerve ending in the skin, where they give rise to different neural crest derivatives and self-renewing cells before birth(Bronner and Simoes-Costa, 2016; Gresset et al., 2015; Morrison et al., 1999)(Fig. 5a); those self-renewing cells, snNCSCs, mature in the sciatic nerve and eventually migrate to the DRG in adult(Fig. 5b).

**Fig. 5.**
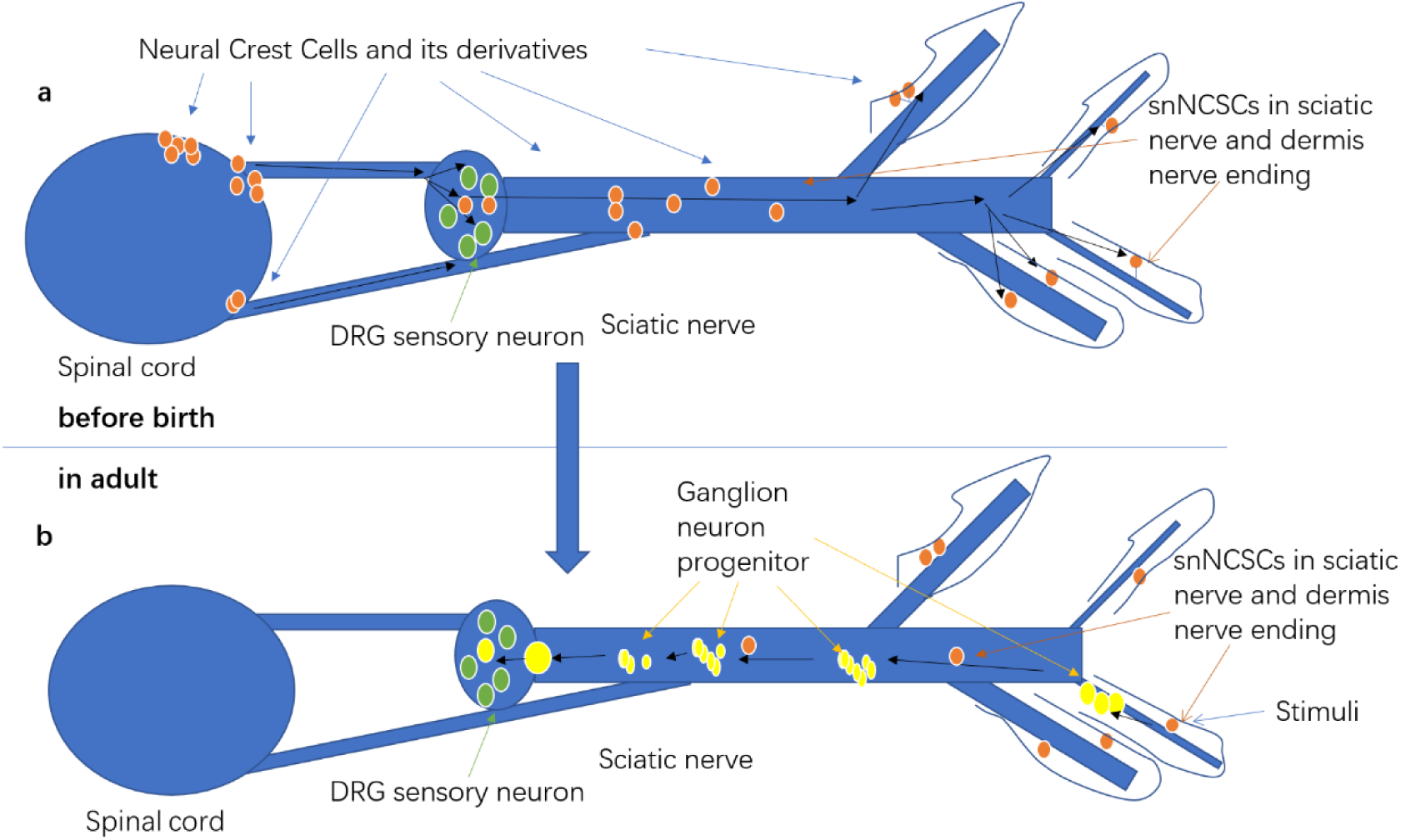
Hypothesis of adult neurogenesis in the sciatic nerve. **a**. Neural crest cells migrate to the DRG and skin along the sciatic nerve before birth. **b**. snNCSCs in the dermis nerve ending and sciatic nerve mature in the sciatic nerve and migrate to the DRG in adults.

In the crush model (Fig. 1a), the sciatic nerve presents a snapshot of adult neurogenesis process outlined in our hypothesis. Along the sciatic nerve from a location distal to the DRG, snNCSCs arranged in a segmental chain of cell groups develop larger cell bodies and undergo a reduction in number. Those segmental form and migration pattern are with same character as the embryonic neural crest(Szabo and Mayor, 2018).

In Nestin-CreER^T2^ rats, those segmental chain comprise neuronal stem cells, progenitors and neurons. Most of those stem cells and progenitors are in quiescence and gradually development along the sciatic nerve.

In Nestin-Cre transgenic rats labeled with the virus via local sciatic nerve injection, we found labeled neurons in the DRG. Without finding of nestin^+^ progenitor cells co-labeled with traced marker in the DRG of those virus injected rats further confirmed our hypothesis that sciatic nerve neurons migrate to the DRG in adult.

Our findings raise two major questions that require further study. First, we must determine the source of snNCSCs. NCSCs in fetal sciatic nerve were shown to have self-renewing ability in vivo and to maintain multipotency in vitro(Morrison et al., 1999). Before birth, boundary cup neural crest stem cell derivatives with multipotency migrate from the neural tube into the nerve roots of skin and the DRG(Gresset et al., 2015). Given the diversity of neural crest cell derivatives(Baggiolini et al., 2015), it would be difficult to determine the source of adult snNCSCs without new methods to track the trajectories of their development.

The second important question is what triggers the stem cells to differentiate. In postnatal parasympathetic ganglia of the head or gut, differentiation of Schwann cell precursors(SCPs) is a programmed organizational process that occurs at a defined time and place(Dyachuk et al., 2014; Espinosa-Medina et al., 2014; Uesaka et al., 2015). In adults, neural crest-derived cells show injury and stress responses, likely involving dedifferentiation and in vivo reprogramming to acquire a new cell fate(Parfejevs et al., 2018). In our 3-day sciatic nerve crush model, the rarity of NC-derived cells in some sciatic nerves indicated that injury and stress may not be the reason why those cells differentiate. On the basis of the observed unpredictable existence and location of those cells in adult sciatic nerve, the differentiation of snNCSCs does not appear to be a programmed organizational process that occurs during specific periods of adulthood.

Although it must be regarded cautiously until verified, we hypothesize that neuronal stem cells in the sciatic nerve are derived from self-renewing cells, which themselves are derived from the neural crest and persist in dermis nerve endings or a branch of the sciatic nerve. The maturation process of those cells is triggered by changes in the environment. These changes are similar to those brought about by an immunogen during the secondary immune response. Once an environmental change (such as the arrival of an immunogen) stimulates the dermis nerve ending or the branch of the sciatic nerve, snNCSCs are activated and stored in the trunk of the sciatic nerve. As mature neurons and snNCSCs in quiescent condition are both in the adult sciatic nerve as EdU staining experiment and single-cell sequencing indicated, the same environmental change will initiate a rapid sensory response. This hypothesis may help explain mammalian responses to environmental change and even drive a new understanding of the evolution of the mammalian sensory system.

Adult neural stem cells are attracting increased interest as potential candidates for cell transplantation therapy for nerve trauma and disease because they are present in tissue that can be harvested from the patient(Parfejevs et al., 2018; Radomska and Topilko, 2017). Moreover, skin stem cells contribute to skin regeneration and wound repair through cellular programs that can be hijacked by cancer cells(Ge and Fuchs, 2018; Mantyh, 2006; Nakada et al., 2011) Our snNCSCs migration model may therefore provide clues to cancer cell migration along the sciatic nerve, expanding knowledge about their role in hijacking the hematopoietic system via blood vessels and lymphatic vessels (Crane et al., 2017). In the future, we hope this work will also facilitate transplantation of adult neuronal stem cells in the sciatic nerve using a method that simulates typical adult sensory reconstruction processes so that we can eventually realize functional sensory reconstruction. Furthermore, our observation of newly born sensory neurons may help elucidate the mechanisms of pain, touch, and other senses, and may one day enable adult sensory reconstruction and help overcome barriers to limb reconstruction.

## Methods

### Animals

Three rat lines were used for this study: Sprague-Dawley (SD), a transgenic Nestin reporter SD rat line that expresses Cre from the endogenous H11 locus, a transgenic Nestin reporter SD rat line that expresses CreER^T2^ from the endogenous H11 locus and pCAG-loxP-3XSTOP-loxP-Tdtomato-WPRE-bGHpA from the endogenous ROSA26 locus. Male experimental and control rats were littermates housed together before the experiment. We produced Nestin-Cre and Nestin-CreER^T2^ knock-in rats via the CRISPR/Cas9 system. First, a single guide RNA (sgRNA) targeting the H11 locus, the SD Nestin promoter-Cre-PA-Nestin Enhancer fragment, was inserted into the H11 locus of rats using CRISPR/Cas9 technology. The rat H11 locus (which is positioned between the Eif4enif1 and Drg1 genes) is ubiquitous, allowing the use of an exogenous promoter to drive higher expression when inserted at the locus. Second, Cas9, sgRNA, and the donor vector were co-injected into zygotes. We transferred the injected zygotes into the oviduct of pseudopregnant SD females. F0 rats were birthed 21–23 days after transplantation, and were identified by PCR and sequencing of tail DNA. Positive F0 rats were genotyped. Lastly, we crossbred positive F0 rats with SD rats to generate heterozygous rats. All animal procedures were performed in accordance with Institutional Animal Care guideline of Nantong University, and were ethically approved by the Administration Committee of Experimental Animals, Jiangsu Province, China.

### AAV Constructs

Recombinant AAV2/9-hEF1a-GFP, AAV2/9-hSYN-GFP, AAV2/PHP.S-hEF1a-DIO-mCherry, and AAV2/PHP.S-hSYN-DIO-GFP vectors were packaged by co-transfection of HEK293 with AAV9 or PHP.S capsid plasmid, helper plasmid, and the corresponding shuttle plasmid(Shanghai Taitool Bioscience Co. Ltd). Virus was collected from each specimen 3 days after transfection and purified with iodixanol discontinuous density ultracentrifugation(Shanghai Taitool Bioscience Co. Ltd). The buffer viral solution was exchanged with phosphate-buffered saline (PBS) plus 5% glycerol using an Amicon ultra-15 spin ultrafiltration (Millipore). Genome copies of final viral solutions were determined by qPCR using primers detecting WPRE and the shuttle plasmid as a standard. The virus titers were approximately 2.50E+13.

### ScaleS and Immunohistology

The transparency procedure was the same as described in other studies that used ScaleS. Briefly, the epineural sheath must first be peeled away from the sciatic nerve sample. The permeability of a sample was enhanced by incubation for 12 h in ScaleS0 solution (20% sorbitol, 5% glycerol, 1 mM methyl-β-cyclodextrin, 1 mM γ-cyclodextrin, 1% N-acetyl-L-hydroxyproline, and 3% DMSO). Second, the permeable (adapted) sample was incubated sequentially in ScaleA2 (10% glycerol, 4 M urea, 0.1% Triton X-100 for 36 hr), ScaleB4(0) (8 M urea for 24 hr), and ScaleA2 (for 12 hr) for permeabilization/clearing. These urea-containing and salt-free ScaleS solutions gradually clear the sample. Then, after descaling with PBS(–) wash for at least 6 hr, the sample was incubated for 36 hours with a fluorescence-labeled primary antibody (Ab) (direct IHC) or a primary Ab and then a fluorescence-labeled secondary Ab (indirect IHC) in an AbScale solution (PBS[–] solution containing 0.33 M urea and 0.1–0.5% Triton X-100). Before refixation with 4% PFA, we applied an AbScale rinse solution to the sample twice, for 2 h each time (0.1× PBS[–] solution containing 2.5% BSA, 0.05% [w/v] Tween-20). Finally, the immunostained sample was optically cleared by incubation in ScaleS4 for more than 16 h (40% sorbitol, 10% glycerol, 4 M urea, and 0.2% Triton X-100). The following antibodies were used: Stmn2 rabbit 1:200 (ProteinTech 10586-1-AP), NeuN mouse 1:200 (Millipore [clone GA5] MAB377), GFP chicken 1:200 (Abcam ab13970), peripherin chicken 1:200 (Aves PER), nestin chicken 1:500 (Aves NES), nestin mouse 1:500 (Chemicon MAB353), CGRP goat 1:200 (Abcam ab36001), plexin C1 mouse 1:500 (R&D Systems AF5375), SSEA-1 mouse 1:200 (Millipore [clone MC-480] MAB4301), mCherry chicken 1:200 (Novus NBP2-25158). All the fluorescence-labeled secondary antibodies were purchased from Invitrogen (1:400).

### EdU and tamoxifen injection and labeling

A stock solution of 10 mg/ml EdU (Invitrogen, A10044) was prepared in normal saline solution (0.9%). A stock solution of 55 mg/ml tamoxifen(Sigma, T5648) was prepared in a 5:1 solution of corn oil:ethanol at 37° in water bath kettle with occasional vortexing overnight. EdU (10 mg/kg) was injected in Nestin-CreER^T2^ rats(6-week old, 200g) with tamoxifen injection(55mg/kg) at the time points showed in the Fig3a. After secondary antibody staining, EdU staining was performed according to manufacturer’s guidelines (Click-iT EdU Alexa Fluor 647 Imaging Kit, Invitrogen).

### Rat Sciatic Nerve Culture Preparation and Treatment

We cultured sciatic nerves from adult 6- to 8-week-old SD rats in 10 cm plates (Corning) coated with poly-D-lysine (Sigma) and laminin (Sigma). The sciatic nerve was cut from below the DRG (omitting all DRG tissue) to the nerve ending. We carefully peeled away the epineural sheath in cold PBS. The collected sciatic nerves were plated as a line at a density of approximately eight sciatic nerves per dish and kept for 20 min at 37°C (make sure it fixed on the plates), and a Neurobasal medium (Invitrogen) supplemented with 2% (vol/vol) B27 (Invitrogen) and 25 ng/mL nerve growth factor (Sigma) was added. Cultured sciatic nerves were maintained for 1 day prior to injection. The culture medium was discarded before the injection. The virus (20 μl/8 nerves, with virus titers of approximately 2.50E+13) was dropped slowly and uniformly onto the sciatic nerve and then incubated for 2 h before adding the culture medium(carefully ensuring that the sciatic nerve did not dry out). Then, the sciatic nerve was cultured for 1–2 weeks before observation to allow for GFP expression from the sciatic nerve, where the cell bodies were placed. For more than one month cultures, the medium was changed every 5 days. We examined neurons using a fluorescence microscope.

### Animal Surgery

For the sciatic nerve lesion experiment, the adult rats (8–10 weeks old) were anesthetized by intraperitoneal injection with 85 mg trichloroacetaldehyde monohydrate, 42 mg magnesium sulfate, and 17 mg sodium pentobarbital. We exposed sciatic nerve at the sciatic notch by making a small incision. The nerve was then crushed at the same position for 30s under the same pressure by a ultra-fine hemostatic forceps, and the crush site was marked with a size 10-0 nylon epineural suture. For the control rats (8–10 weeks old), the sciatic nerve was exposed but left uninjured. After surgery, the wound was closed, and the rats were allowed to recover for two hour.

For the sciatic nerve AAV injection, we anesthetized the adult rats (6–8 weeks old, normal rats and 6 weeks old, Nestin-Cre rats) with an intraperitoneal injection of complex narcotics (85 mg trichloroacetaldehyde monohydrate, 42 mg magnesium sulfate, and 17 mg sodium pentobarbital) and carefully opened the skin and muscle and to expose the sciatic nerve. Then 10 cm capillary glass tubes (Sutter Instrument, Novato, CA) were pulled using a micropipette puller (model 720, David KOPF Instruments, Tujunga, CA). The tips of the pulled tubes were pinched with forceps to create pipettes with an external diameter of approximately 10 μm. A 2.5 μl volume of AAV2/9 (virus titers were approximately 2.50E+13) was gradually injected into the sciatic nerve with one pump of a microsyringe pump at a rate of 1 μl/min (Stoelting Instruments). The needle tip was inserted into the epineural sheath, and the drops caused it to plump up. After three injections at three different sites along the sciatic nerve, the wound was closed, and the rats were allowed to recover for two hour.

For the tail AAV injection, we placed the 3-week-old rats in a restraint device. The tail was stabilized between the investigator’s thumb and forefinger. To soften the skin, the tail was prepared in 40°C water for 5 min and then sterilized by 70% ethanol. The injection started at the distal part of the tail with an insulin syringe. With the tail under tension, the needle was inserted approximately parallel to the vein at a depth of at least 3 mm. A 30 μl solution of AAV-PHP.S (the virus titers were approximately 2.50E+13) mixed into 150 μl PBS was slowly injected over 3 min. After the vein blanched, the needle was kept in position for 1 min. The rats were allowed to recover for two hour and then returned to their home cages.

### Preparation of Individual Cells From Adult Rat Sciatic Nerve

We conducted single cell sequencing experiments on SD rats subjected to hSYN-GFP AAV2/9 and AAV2/9-hEF1a-GFP virus sciatic nerve injection in vivo and in vitro. The rats were euthanized by cervical dislocation, and the sciatic nerve was immediately immersed in ice-cold Dulbecco’s Phosphate-Buffered Saline (DPBS, Corning). The dissected sciatic nerve was cut from the distal end of the DRG to the end of the sciatic nerve (leaving out the DRG) and we carefully peeled away the epineural sheath in cold PBS. Collected sciatic nerves were cut into pieces under a fluorescence microscope. The GFP^+^ piece was collected and incubated in Hibernate A (BrainBits) containing papain (100 U; Sigma) at 37°C for 2 h with intermittent flicking. After removing enzymes, the collected pieces were trypsinized for 20 min at 37°C. The tissue was triturated into an individual cell suspension using a 1 ml pipette. We removed the trypsase and cellular debris with three rounds of mild centrifugation at 1000 ×g and a Hibernate A minus Ca^2+^ and Mg^2+^ wash (BrainBits). The individual cell suspension was plated into a glass-bottom plate and collected using glass pipettes under a fluorescence microscope. The glass tip was broken off and left in each PCR tube containing lysis buffer (Vazyme Biotech) with water (2.4 μl), RNase-free DNase (0.2 μl) and murine origin RNase inhibitor (0.25 μl).

### Library Preparation, Clustering and Sequencing

We used the Vazyme method, followed by cDNA amplification as described below. Whole transcriptome amplification was performed using the Discover-scTM WTA Kit V2 (Vazyme, N711). First, 124 active cells were isolated and transferred into a lysis buffer. Then, mRNA was copied into first-strand cDNA using Discover-sc Reverse Transcriptase and oligo dT primer. At the same time, we added a special adapter sequence to the 3’ end of the first-strand cDNA. Full-length cDNA enrichment was performed by PCR, and the products were purified by VAHTSTM DNA Clean Beads (Vazyme, N411). Next, we performed quality control using the WTA cDNA. The cDNA concentration was measured using a Qubit DNA Assay Kit in a Qubit 3.0 Fluorometer (Life Technologies, CA, USA). DNA fragment size was tested using an Agilent Bioanalyzer 2100 system (Agilent Technologies, CA, USA). A total of 1 ng of qualified WTA cDNA product per sample was used as input material for the library preparation.

We generated sequencing libraries using the TruePrep DNA Library Prep Kit V2 for Illumina (Vazyme, TD503), following the manufacturer’s recommendations. First, cDNA was randomly fragmented by the Tn5 transposome at 55°C for 10 min at the same time as a sequencing adapter was added to the 3’ adenosine on the fragment. After tagmentation, the stop buffer was added directly into the reaction to end tagmentation. PCR was performed, and the products were purified with VAHTSTM DNA Clean Beads (Vazyme, N411We conducted preliminary quantification of the library concentration using a Qubit DNA Assay Kit in Qubit 3.0. Insert size was assessed using the Agilent Bioanalyzer 2100 system, and if the insert size was consistent with expectations, it was more accurately quantified using qPCR with the Step One Plus Real-Time PCR system (ABI, USA).

We identified the neuron-like cells via Stmn2 expression. We identified 94 Stmn2^+^ cells in vitro and six Stmn2^+^ cells in vivo using qPCR before sequencing. 14 Stmn2^-^ cells were identified as negative control. 10 DRG neurons were identified as positive control.

Clustering of the index-coded samples was performed on a cBot Cluster Generation System (Illumina) according to the manufacturer’s instructions. After cluster generation, the library preparations were sequenced on an Illumina Hiseq X Ten platform with a 150 bp paired-end module.

### Bioinformatic Analysis

Samples were then normalized by down sampling to a minimum number of 124 transcripts per cell for the clustering analyses or a minimum of 100 transcripts per cell for differential gene expression analyses. Cells with fewer transcripts were excluded from the analyses. The modularity optimization technique SLM was used for unsupervised cell clustering. We used t-SNE to place cells with similar local neighborhoods in high-dimensional space together(McDavid et al., 2013).

## Data availability

The 124 single-cell sequencing data(sample gene expression in FPKM) is available on Dryad (https://doi.org/10.5061/dryad.xgxd254f6) for the bioinformatic analysis.

## Acknowledgements

I would like to thank H. Wang for providing the Nestin-Cre rats; F. Liu for helping with microscopes; X. Li for helping with the imaging of sciatic nerve; S. Zhou, l. Zhao, J. Qin, P. Li, J. Li, P. R. Williams for helping with sciatic nerve crush and injections; C. Zhou for helping with histology; Z. Wei and C. Li for helping with single-cell sequencing. L. Cai for helping with the design of the sketch map. This study is grateful for support from National Natural Science Foundation of China (Grant No. 31730031). I apologize that all relevant publications could not be cited.

